# Directed Micro-Ecology Application System: On-site One-step Fermentation Facilitating Microbial Application Through Theoretical and Equipment Innovation

**DOI:** 10.1101/2024.06.15.599119

**Authors:** Rui-qiang Ma, Yun-feng Luo, Wei-min Zhao, Jian-feng Guo, Yanjie Li, Wen-jing Cui

## Abstract

Functional microbial agents play a crucial role in various fields such as agriculture, feed fermentation, aquaculture, and environmental protection. However, traditional microbial agents were confronted with critical challenges such as limited shelf-life, reduced activity, and inconsistent efficacy. In this case, we innovatively proposed the concept of Directed Micro-Ecology (DME) and developed its application system, including a core module named DME intelligent fermentor (DME25). Over 40 functional strains, including bacterial strains and fungus strains, were successfully cultured to 10∼50 ×10^8^ CFU/mL within 20∼48 h and maintained a relatively low contamination rate (<2.5%). Finally, the stability and effectiveness of these DME-fermented strains were validated in different application areas, all of which exhibited perfect functional characteristics. Firstly, the bacillus strains inhibited the progression of wilt disease and significantly improved the growth of tomatoes. Secondly, all tested lactobacillus strains improved the nutrition and quality of fermented feed, complying with feed industry standards. Lastly, the ammonia nitrogen concentration, nitrite concentration of aquaculture water and phosphate concentration, COD of aquaculture tail water were significantly reduced within 1∼4 d. The successful application of the DME intelligent fermentor in different fields marks a pivotal breakthrough in technological innovation of microbial agents on-site one-step fermentation. This technological advancement opens new avenues for enhancing the stability and effectiveness of microbial agents, infusing powerful impetus to the development of microbial application.

## Introduction

Microbial agents were extensively used in multiple fields based on their significant biological functions. Increasing numbers of farmers and agricultural enterprises are adopting microbial agents to replace chemical pesticides, achieving biological control and increasing crop yields (Gouda et al., 2018; Huang et al., 2020; Jiao et al., 2021). Simultaneously, microbial agents are widely used in projects such as sewage treatment and soil remediation, accelerating the degradation of organic waste and reducing environmental pollution (Hasr Moradi Kargar and Hadizadeh Shirazi, 2020; Ilyin et al., 2004; Kim and Cho, 2022; Zhang et al., 2023). In industrial production, particularly in the preparation of bioenergy, microbial agents are expected to replace traditional fossil energy production methods, reducing adverse environmental impacts (Matsushika et al., 2009; Muras et al., 2021). In the livestock industry and aquaculture, their role in improving feed utilization, enhancing animal immunity, improving environmental hygiene, and promoting sustainable livestock development will provide important support for industry healthy development (Shao et al., 2021). Therefore, microbial agents play a vital role in achieving sustainable development, promoting ecological balance, protecting the environment, and improving human quality of life (Pourfadakari et al., 2021). As the pursuit of sustainable development and environmental protection awareness increases, the prospects for their application will become even broader (Adedayo et al., 2022; Meng et al., 2022).

Currently used microbial strains mainly include Gram-negative strains (*Sinorrhizobium, Pseudomonas, Nitrobacter*), Gram-positive strains (*Bacillus, Lactobacillus, Paenibacillus*) and fungus strains ( *Trichoderma, Saccharomyces*) (Dobrzyński et al., 2023; Sadare and Daramola, 2023; Song et al., 2022; Tyśkiewicz et al., 2022; Woo et al., 2023). In general, bacillus strains were not sensitive to various adverse stresses and were widely applied, due to their ability to produce spores (Jeżewska-Frąckowiak et al., 2018; Paul et al., 2019; Todorov et al., 2022). However, other non-spore-forming microbial strains were hyper-sensitive stress tolerance, resulting in shorter shelf-life of their products, unstable application effects, and poor adaptability in the environment, leading to ineffective survival and colonization (Du et al., 2021; Schumpp and Deakin, 2010; Tittabutr et al., 2007). Therefore, improving the activity and quality stability of microbial agent products was crucial for maintaining their stable and efficient application (Chen et al., 2021; Poole et al., 2018). In recent years, increasing numbers of researchers have been attempting to overcome the shelf-life limitations of microbial agents. Nowadays, there are three common methods, including adding protective agents to liquid formulations, spray drying or freeze-drying to produce microbial powder and on-site fermentation through industrial small-scale fermenters, were extensively adopted (Biradar, 2018; Elsakhawy et al., 2021; Perry, 1995). Firstly, adding protective agents was the simplest way with a low cost and short period (Bellali et al., 2020), but the extension of the shelf-life achieved through this method was limited, and the high transportation costs of liquid formulations also limited its development. Secondly, the spray drying technical process was mainly suitable for spore-producing bacteria (Adjallé et al., 2011; Broeckx et al., 2016; Dianawati et al., 2016) and the finished spore powder faces the risk of low spore germination rates during use. Although freeze-drying could effectively extend the shelf-life of non-spore-forming strains (Yang et al., 2023), its high production cost resulted in high usage costs. Finally, using industrial fermentors for on-site fermentation exhibited great enormous advantage of standardized pure culture cultivation (Nobre et al., 2018; Wang et al., 2019). Despite that, the high equipment procurement cost, complex operation, and requirement for specialized fermentation personnel were the three stubborn barriers for customers. To date, there’s still no efficient and straightforward solution to solve the short shelf-life and maintain the activity of microbial agents. Interestingly, probiotics (*Lactobacillus*) were widely used for homemade milk fermentation, because of their role in gut microbiome modulation and gastrointestinal health benefits (Vasudha et al., 2023). Besides, *Rhodopseudomonas palustris*, attributing to its capacity of autotrophic photosynthesis, was commonly cultured in outdoor, closed transparent water tanks by aquaculture clients (Carlozzi et al., 2006; Harwood, 2022). Collectively, immediate use after fermentation may shed light on alternative solutions to solve the problem of the shelf-life of microbial and its products easily.

Generally, the misconception of using pure microorganisms for applications was commonly existing and unsuspicious. Actually, microbial agents were used in fields such as feed fermentation, organic material maturation, soil improvement and remediation, and sewage purification under open ambient conditions (Cui et al., 2023; Hong et al., 2022; Stelma, 2018). It seems clear that, from a microbial functional standpoint, as long as the core population comprises the desired functional microorganisms, equivalent functionality can be attained. This concept bypassing the complex aseptic processes and will accelerate the conversion of functional microorganisms. Based on this concept, Green Nitrogen proposed directed micro-ecology (DME) and developed a DME intelligent fermentor through equipment and fermentation process innovation. In detail, the target functional microorganisms would dominate the final fermentation population under directed selection pressures, including values of fermentation Tm, pH, EC, DO, and selective nutrition, even under simple conditions of pasteurization. This DME intelligent fermentor with rapid proliferation and fermentation of target microorganisms in application scenarios, ensuring the activity of microorganisms, greatly improving product quality, and ensuring the application stability and efficiency of microbial agents. The DME application system was suitable for many fields and may help customers to achieve the desirable application performance of most agricultural microbial agents.

## Materials and Methods

### Strains and Materials

The strains used in this study are shown in Table S1. All DME directional mediums 1∼5 are shown in Table S2. The fermentation yield of these strains can reach 1.5 to 5.9×10^9^ CFU/mL, with a contamination rate controlled below 2.5% (Fig. S1).

The tomato seeds (*Solanum lycopersicum* Miller, *cv*. Zhongza201) were purchased from the Chinese Academy of Agricultural Sciences. The feed materials were provided by Zhongnong Chuangda (Beijing) Environmental Protection Technology Co., Ltd. The aquaculture water and the aquaculture tail water samples were collected from small-scale greenhouse aquaculture ponds for South American white shrimp in Dongying, Shandong Province.

### DME Fermentation of Different Strains

Validate the effects of fermenting different strains using DME fermentation. Firstly, turn on the DME and click on the “feed” button to allow the machine to automatically add water. Next, add the culture medium into the fermentation vessel and put the inoculant into the inoculation chamber. The DME fermentation machine will then start intelligent fermentation. Once the water level reaches the set standard, begin heating for sterilization. Sterilization is conducted at 85∼90°C for 15 min. After sterilization, the equipment will start cooling down until the fermentation temperature (37°C for bacillus strains). At this point, the inoculant in the inoculation chamber will be automatically injected into the fermentation vessel and automatically following the preset program for aeration and agitation during fermentation. When the program is completed, click the “output” button to obtain the usable inoculant.

### Microbial Population and Contamination Rate Calculation

The viable microbial count was determined using the dilution plate method. The microbial suspension was serially diluted from 10^-1^ to 10^-7^, and appropriate dilutions were selected to pipette 100 µL onto corresponding solid plates. The plates were then inverted and incubated at 28°C for 2 d in a constant temperature incubator. Microbial colonies were counted, and the final microbial population and contamination rate were calculated.

### The Biocontrol and Growth-Promoting Test of Tomato

Initially, the seeds were subjected to a disinfection process, which included agitation and immersion in 95% ethanol for 1 min, followed by agitation and immersion in 2.5% NaClO for 3 min. The seeds were then rinsed 8-10 times with sterile water. Subsequently, the pre-treated seeds were placed on damp gauze, covered with an additional layer of gauze, and positioned within a petri dish. The dish was then covered with a breathable membrane to maintain moisture. The seeds were incubated at 25°C for 3 days until they began to germinate. Seeds with uniform sprout lengths were selected for sowing. Following planting, the nutrient soil was thoroughly watered and covered with a film. The seeds were then cultivated in a controlled environment chamber to ensure optimal growth conditions. Each treatment consisted of 40 tomato seedlings.

*Ralstonia solanacearum* was inoculated into NA broth, 28°C, 150 rpm, 24 h. During the 20 d seedling stage, the tomato seedlings were inoculated with *R. solanacearum* at a concentration of 10^8^ CFU/mL, with 0.5 mL per seedling applied to the tomato roots. The following day, each seedling was inoculated with the biocontrol agent 0.5 mL GN125 at a concentration of 5×10^8^ CFU/mL to determine its disease resistance. Additionally, the culture medium and the biocontrol agent GN125 were inoculated separately to assess its growth-promoting effect. Plant height was measured by taking the distance from the soil surface to the top growth point of tomato plants. Additionally, stem diameter at 2 cm above the soil surface was measured. Starting from the bottom, the first fully developed trifoliate leaf was selected for chlorophyll content measurement (SPAD-502PLUS).

### Silage and Corn Feed Experiment Process

The corn was harvested at the early wax ripeness stage and cut into 2 cm sections. Each section was then thoroughly mixed with the respective microbial agent to achieve a final strain concentration of 10^5^∼10^6^ CFU/g. The experiment included three treatment groups: the control group without the addition of microbial agents, and three groups with strains addition. Each treatment had three replicates, with each replicate contained in a 40 kg silage bag. The bags were vacuum-sealed and stored in a cool, dark place for approximately 2 months, with the highest daily temperature averaging 20°C and the lowest averaging 11.3°C. The corn feed underwent a similar fermentation process.

The quality of fermented silage was measured by Bayannur Hemao Muye Co. Ltd. The concentration of aflatoxin and vomitoxin of the corn feed was measured by kit (BA0141, Shanghai Youlong Biotech Co,.Ltd) according to its manufacture.

### Aquaculture Water and Aquaculture Tail Water Management and Parameter Analysis

Tested strains were respectively added to containers with 300 mL of water samples, ensuring a bacterial concentration of 1×10^6^ CFU/mL in the water. The containers were then placed in a water bath and incubated at 28°C with aeration, equaling one volume of the water. The concentrations of ammonia nitrogen and nitrite in the aquaculture water and the concentrations of phosphate and COD in the aquaculture tail water were measured 1∼4 d.

Multi-parameter water quality analyzer was purchased from Xiamen PanTian Biotech Co., Ltd. The determination of ammonia nitrogen was conducted using the Nessler reagent colorimetric method, while the determination of nitrite was carried out using the α-naphthylamine colorimetric method. The concentration of phosphate was measured by using the molybdenum blue method. COD was measured by COD on-line Monitor (Shanghai BoQu Instrument Co., Ltd, CODG-3000) according to its manufacture.

## Data Analysis

Graphs for experimental data were generated using GraphPad software. Analysis of variance (ANOVA) single-factor variance analysis and t-tests were conducted using SPSS software.

## Results

### 1. Basic Principle and Application Development System of Directed Micro-Ecology (DME)

To achieve on-site standardized production and convenient operation of microbial products, Green Nitrogen Biotech Co., Ltd. (referred to as “NBio”) proposed the concept of Directed Micro-Ecology (DME) (Fig.1a). The principle was to apply directional selective pressures, such as the culture medium, fermentation time, temperature, EC value, rotation speed, and dissolved oxygen level, to create a microbial ecosystem that promotes the growth of the desired functional microorganisms. This ecosystem should contain both the target microorganisms and limited environmental microorganisms, to maintain the dominance of the target functional microorganisms. This pressure enables the target microorganisms to gain an advantage in inter-species competition, and occupy a dominant position in the microbial community within a specific cultivation period, manifesting as dominant microorganisms. Moreover, the low contamination rate from environmental microorganisms ensures that the target microorganisms perform effectively in practical applications. Based on the DME principle, NBio developed an application development system consisting of four major modules to establish the ‘FNPP’ cycle, achieving the iterative upgrading of a DME application system (Fig.1b). This system identifies target functional microorganisms (Function, F) based on application requirements, determines corresponding directed culture medium (Nutrition, N) and fermentation process models (Process, P), and accomplishing on-site standardized fermentation and application of target functional microorganisms through a DME intelligent fermentor (Production, P). By introducing the corresponding culture medium and inoculant into the DME intelligent fermentor, an integrated and intelligent fermentation process can be initiated (Fig.1c).

**Fig. 1.**
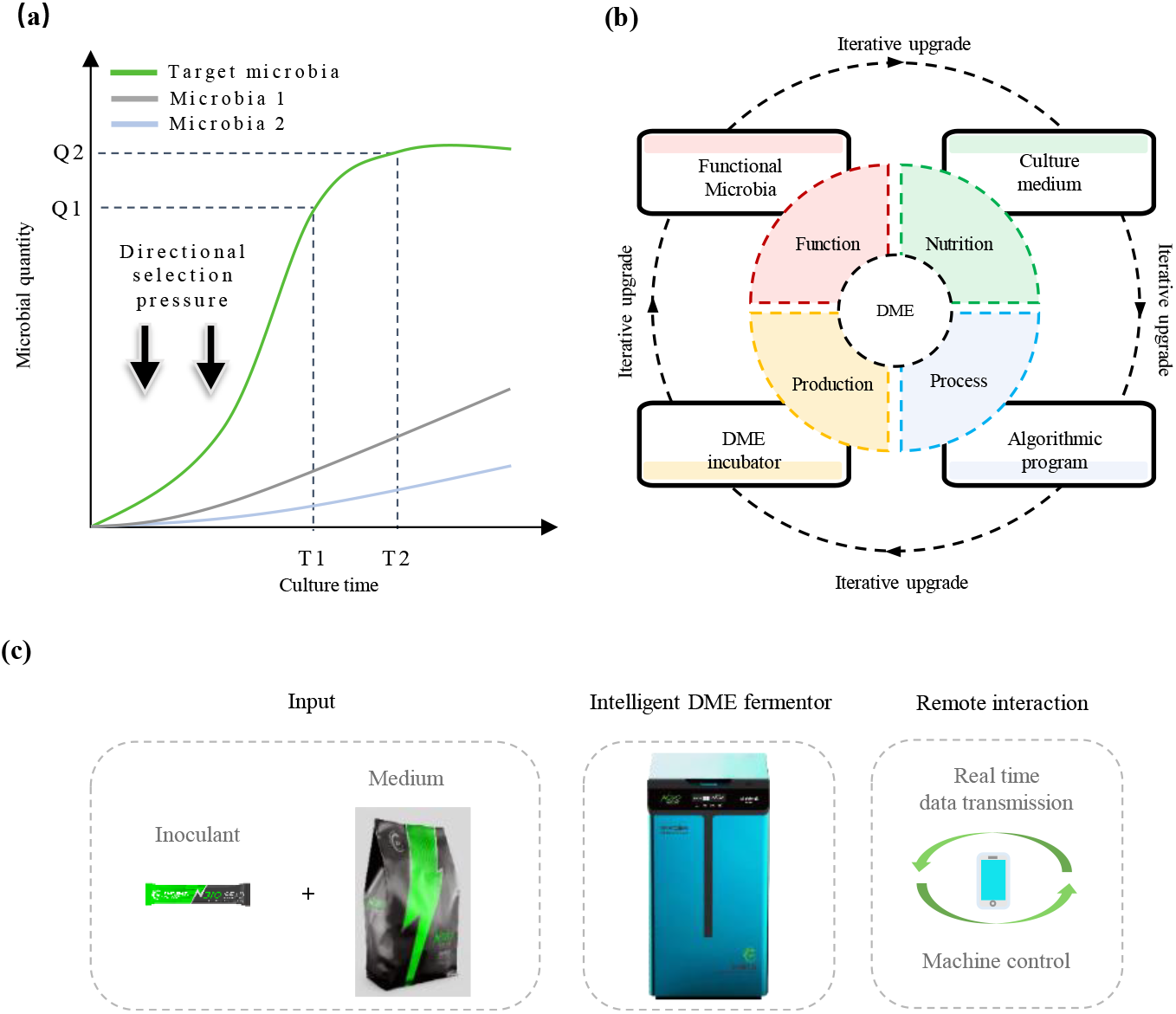
Directed Micro-Ecology (DME) and its application development system. a, Basic principle of Directed Micro-Ecology (DME). The directional selection pressure was applied to favor the dominance of the desired population (Q1-Q2) of functional microorganisms within a specified cultivation time (T1-T2); b, The ‘FNPP’ development cycle, FNPP standing for Function-Nutrition-Process-Production; c, Intelligent DME incubator (DME25) and fermentation consumables, including inoculant and culture medium.

The DME intelligent fermentor (DME-25), which includes temperature control components, heating and cooling modules, a stirring motor, an air pump, a filtration membrane, and an automatic inoculation chamber. It operates within a temperature control range of RT to 95°C, with adjustable parameters for temperature, rotation speed, and aeration volume. The main parameters are summarized in Table 2. This system can be operated by individuals without specialized training and does not need to be monitored after startup, resulting in significant savings on labor costs. Additionally, DME-25 was equipped with sensors that allow for real-time monitoring and recording, meeting research needs. This feature enables precise control and documentation of the fermentation process, making the DME-25 an ideal tool for both practical applications and academic research.

**Table 1.**
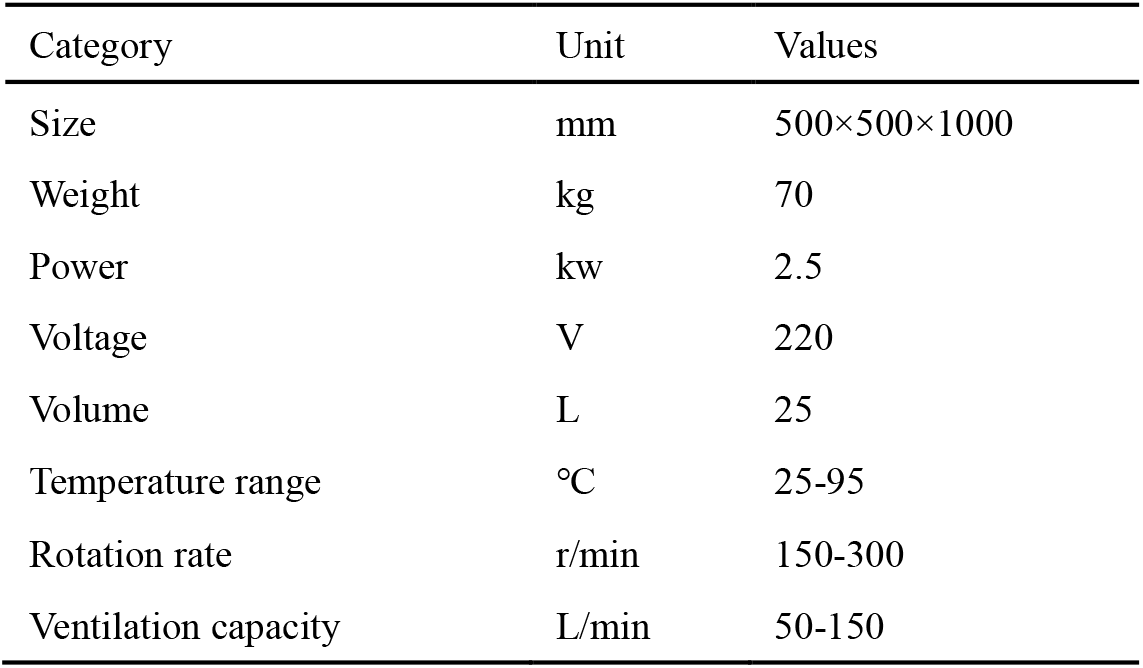
Parameter of the DME-25 fermentor.

**Table 2.**
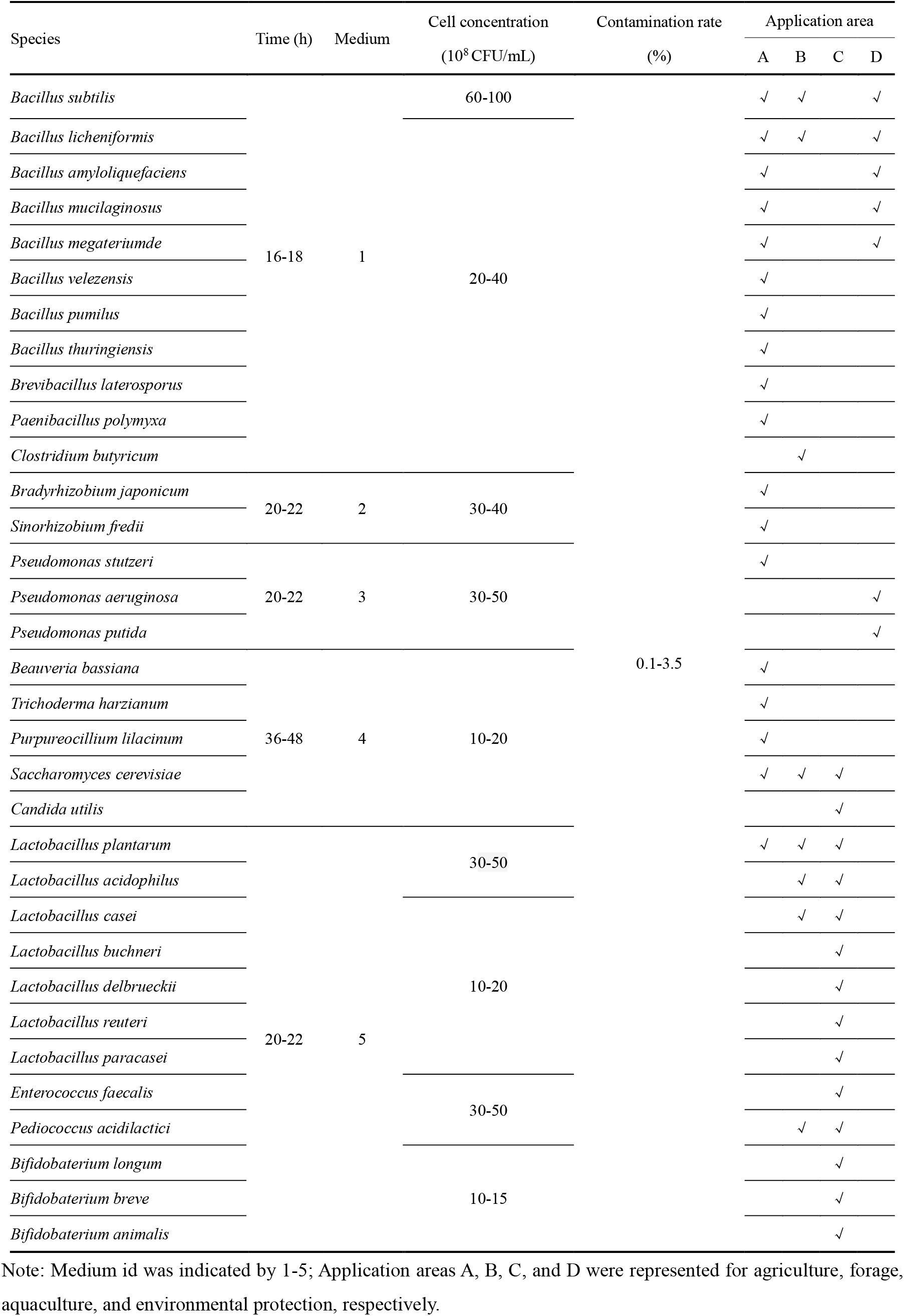
DME incubator partial matched strains.

### 2. Fermentation Tests of Commonly Used Microbial Agents on DME Fermentor

We conducted DME fermentation tests on common microbial strains used in agriculture, aquaculture, forage production, and environmental protection (Table 2). The results showed that Bacillus, when adapted to medium 1, achieved a bacterial count of 20-100×10^8^ CFU/mL after 16-18 hours of fermentation in the DME fermentor. When adapted to medium 2, rhizobium strains reached a count of 30-40×108 CFU/mL after 20-22 hours of DME cultivation. When adapted to medium 3, *Pseudomonas* achieved a count of 30-50×10^8^ CFU/mL after 20-22 hours of cultivation. *Lactobacillus, Staphylococcus*, and *Lactobacillus*, when adapted to medium 5, reached a count of 10-50×10^8^ CFU/mL after 20-22 hours of cultivation. For several fungal strains adapted to medium 4, the required fermentation time was 36-48 hours, slightly longer than for bacterial fermentation, with a count of 10-20×10^8^ CFU/mL. Excitingly, the contamination rate for all tested strains was controlled below 3.5% ( Table 2 and Fig. S1). These results demonstrate the effectiveness of the DME system in cultivating a wide range of microorganisms under optimal conditions, ensuring high yield and viability across different species.

### 3. The Application Effect of Functional Microorganisms by Using DME Intelligent Fermentor

#### 3.1 Validation of GN125 Efficacy in Tomato Growth-Promoting and Disease Resistance Using DME

##### Fermentor

To validate the efficacy of functional microorganisms expanded using the DME fermentor, we conducted a tomato pot experiment. Data were collected at the seedling stage and harvest stages for verification, and the results were presented in Fig. 2. After inoculation with *Ralstonia solanacearum* and subsequent treatment with GN125, significant increases were observed in plant height, stem diameter, fresh weight, and leaf chlorophyll content compared to plants inoculated solely with *Ralstonia solanacearum*. These findings suggested that *Bacillus subtilis* GN125 fermented by DME exhibits disease-resistance properties. Additionally, growth-promotion assays showed that plants irrigated with *Bacillus subtilis* GN125 had significantly enhanced the plant height, stem diameter, fresh weight, and leaf chlorophyll content compared to untreated controls. These results confirmed that *Bacillus subtilis* GN125 fermented by DME possesses growth-promoting properties. Overall, the DME-fermented *Bacillus subtilis* GN125 maintained its superior disease resistance and growth-promotion properties.

**Fig. 2.**
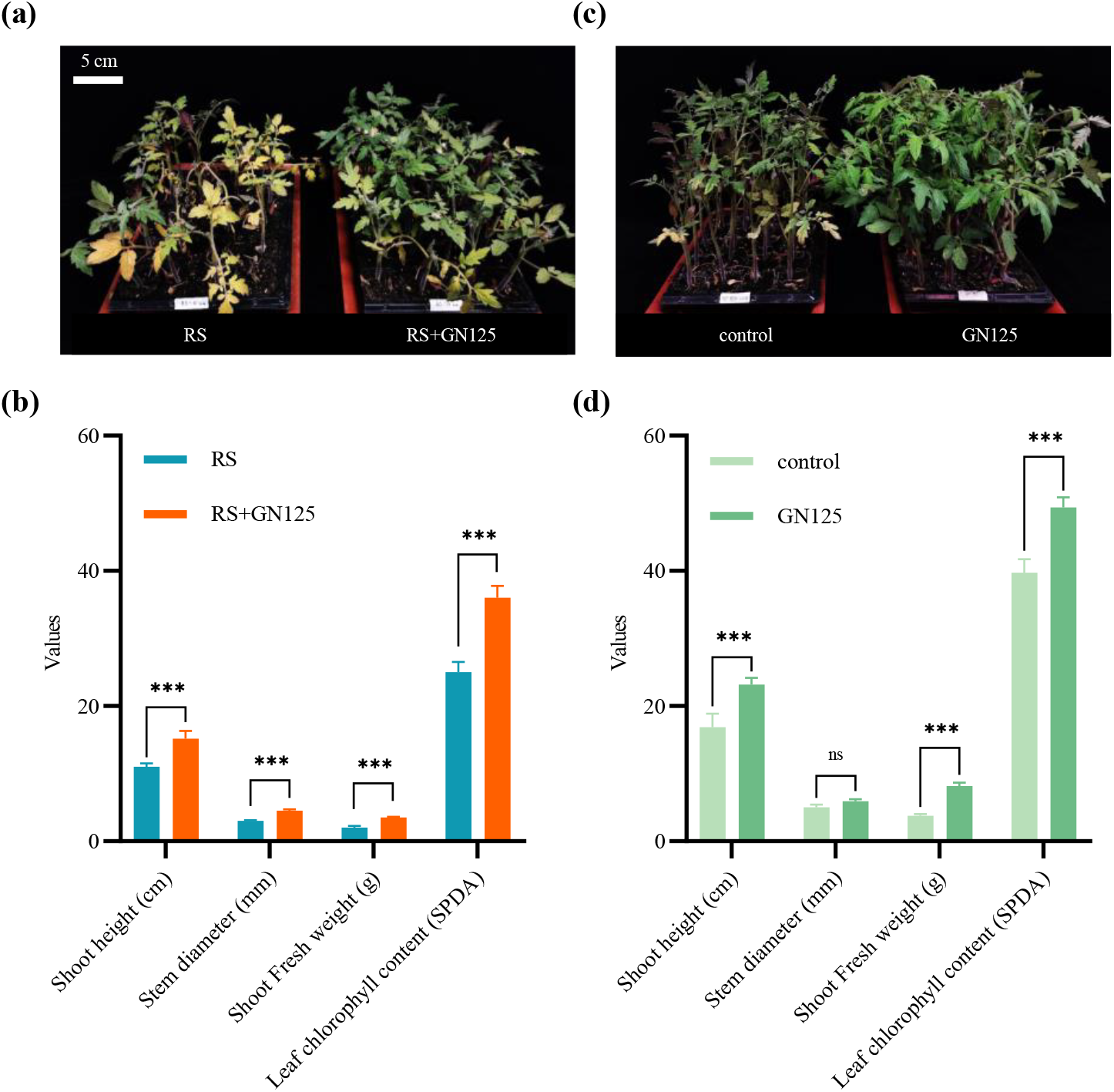
Tomato growth promotion and tomato bacterial diseases biocontrol The phenotype. (a) and the physiological indexes (b) of tomato treated with bacterial wilt pathogen *Ralstonia solanacearum* (RS) and *Bacillus subtilis* GN125. The phenotype (c) and the physiological indexes (d) of tomatoes treated with growth-promoting bacillus strain GN125. Physiological indexes include the shoot height, stem diameter, shoot fresh weight, and leaf chlorophyll content (measurement indicated in bracket). Error bars represent standard deviation (SD). The significance test was accomplished by *t*-test. *, significant *p* value at 0.05 level; ***, significant *p* value at 0.001 level; ns, no significant difference.

#### 3.2 Enhancement of Silage Quality and Nutrition through DME-Fermented Microbial Agents

We evaluated the effects of DME-fermented microbial agents on fermented silage, considering sensory characteristics, quality, and nutritional composition. The results are presented in Table 3. The fermentation treatments using *Lactobacillus plantarum* GN1125, GN251, and GN1367 showed a less uniform color but a stronger acid aroma compared to the control. However, there was no noticeable difference in texture. The pH values of the treated silages were slightly higher, but still within a beneficial range to effectively inhibit harmful microorganisms. The ratio of ammonia nitrogen to total nitrogen remained unchanged, indicating that protein degradation did not increase, thus maintaining feed quality. The dry matter content improved to varying degrees, with GN251 showing the most significant improvement, followed by GN1367 and GN1125 with a 3% increase. The crude protein content increased with GN1125 and GN1367 treatments, while GN251 had no significant effect. Nutritionally, the starch content increased and both neutral detergent fiber and acid detergent fiber contents decreased, resulting in improved feed digestibility and palatability. The fermented silage from all strains met the requirements of the level 1 feed standard, while the unfermented silage only met the level 2 feed standard. In conclusion, the microbial treatments enhanced feed quality, with GN251 demonstrating the most significant improvements.

**Table 3.** The evaluation of fermented silage.

| Evaluation category |  | Control | *Lactobacillus plantarum* |  |  |
| --- | --- | --- | --- | --- | --- |
|  |  |  | GN1125 | GN251 | GN1367 |
|  |  | Scores |  |  |  |
| Sensory (30) | Color | 10<br>(green) | 8<br>(yellow) | 8 (yellow) | 8 (yellow) |
|  | Smell | 8 (good) | 10 (fresh) | 10 (fresh) | 10 (frsh) |
|  | Texture | 10 (soft) | 10 (soft) | 10 (soft) | 10 (soft) |
| Quality (30) | pH | 13 (3.63) | 13 (3.68) | 13 (3.78) | 13 (3.78) |
|  | Ammonia N/Total N (%) | 15 (4%) | 15 (4%) | 15 (4%) | 15 (4%) |
|  | Dry matter/DM (%) | 6 (29%) | 8 (32%) | 8 (38%) | 8 (36%) |
| Nutrient (40) | Crude protein (%) | 8 (8%) | 8 (11%) | 8 (8%) | 8(11%) |
|  | Starch (% DM) | 4 (16%) | 4 (17%) | 8 (33%) | 6 (20%) |
|  | Neutral detergent fiber (% DM) | 0 (60%) | 6 (50%) | 8 (44%) | 6 (48%) |
|  | Acid detergent fiber (% DM) | 4 (28%) | 6 (24%) | 8 (21%) | 8 (22%) |
| Total score |  | 78 | 88 | 96 | 92 |
| Grade |  | 2 | 1 | 1 | 1 |

Note: The fermented silage was evaluated by three indicators, including sensory, quality, and nutritional composition. The maximum score for evaluation was 100 points. 85-100 points, grade 1; 70-84 points, grade 2; 60-69 points, grade 3. Control group represent for unfermented silage. GN1125, GN251, and GN1367 belong to *Lactobacillus plantarum*.

#### 3.3 Application testing of the DME fermentor in fermented corn feed

The efficacy of fermented corn feed with commonly used strains on DME was evaluated (Table 4). The results indicated that, with the exception of *L. buchneri* GN1061, all treatments effectively reduced the pH of the fermented corn feed and inhibited the growth of harmful microorganisms better than the control. Additionally, all four treatments had lower levels of aflatoxin and vomitoxin compared to the control, ensuring the safety of the feed. Furthermore, all treatments including the control group, complied with the standards for fermented feed, with all indexes falling below the safety threshold. It’s noteworthy that the addition of microbial agents did not result in contamination, also ensuring both the quality and efficiency of the fermentation process.

**Table 3.**
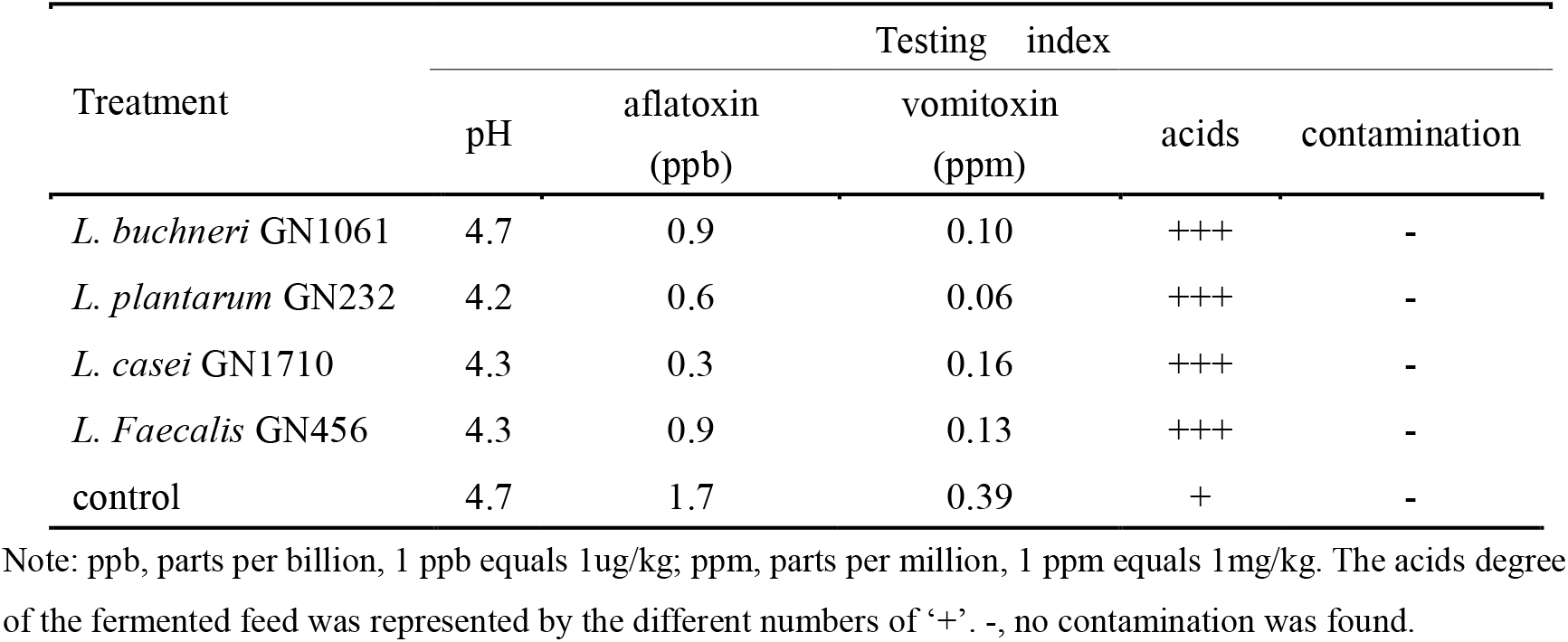
The evaluation of fermented corn feed.

#### 3.4 Enhancing Aquaculture Water and Aquaculture Tail Water Management with DME-Fermented Microbial Agent

Similarly, we conducted the efficacy verification of functional microbial in aquaculture water and aquaculture tail water management. The results showed that after the addition of *B. subtilis* GN2036, the levels of ammonia nitrogen, nitrite, phosphate, and COD in both the aquaculture water and the aquaculture tail water were significantly reduced compared to control (Fig.3 a, b). In contrast, when *L. plantarum* GN104 was added, there was a noticeable reduction in ammonia nitrogen and nitrite levels of aquaculture water, while the reductions in phosphate and COD levels of aquaculture tail water were slight and not as significant (Fig.3 c, d). These findings indicated that the addition of these microbial agents improved the water quality in aquaculture, particularly in reducing the level of ammonia nitrogen, nitrate, phosphate, and COD which was harmful to aquaculture and the environment.

**Fig. 3.**
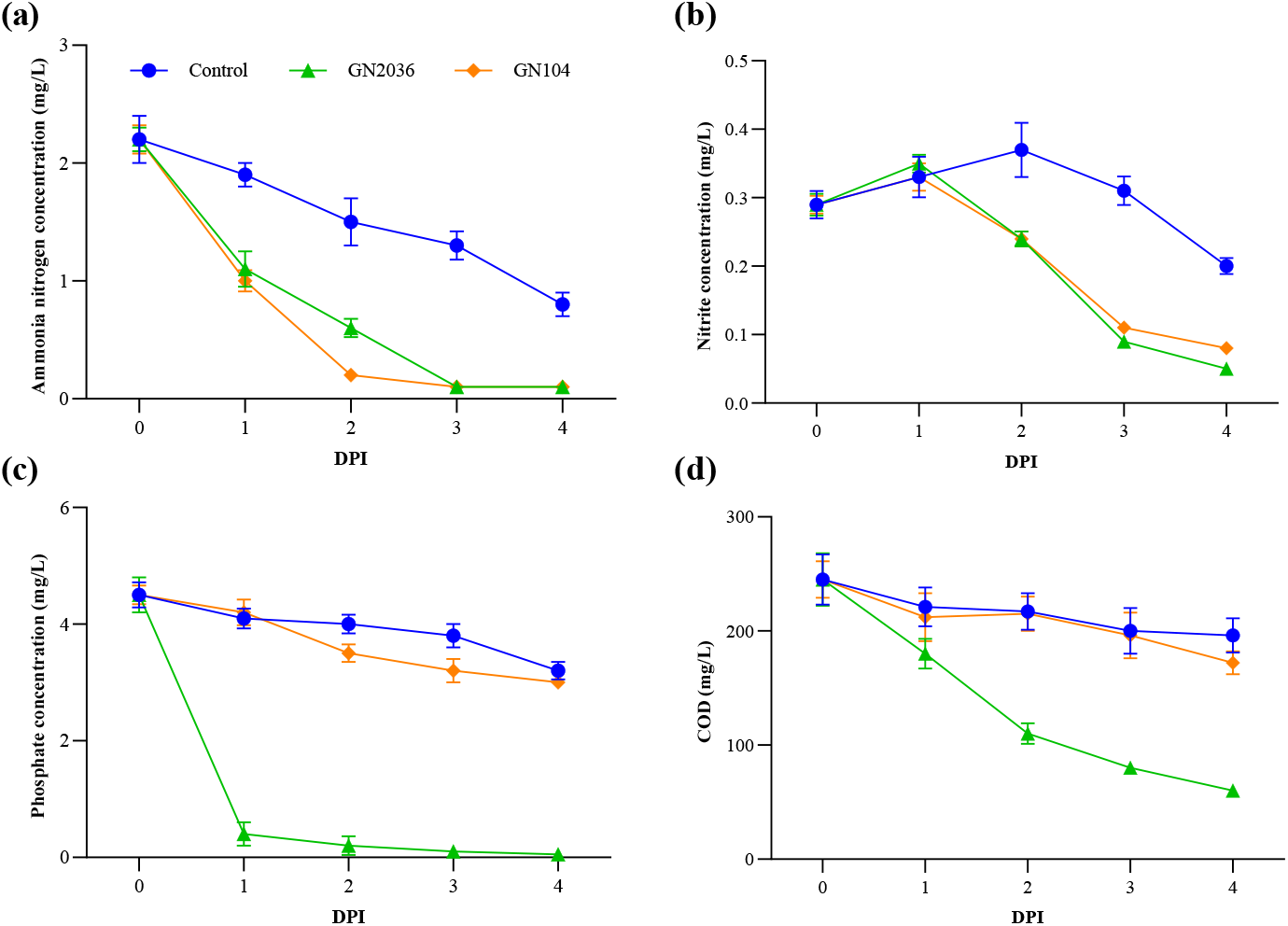
The aquaculture water and aquaculture tail water management. *B. subtilis* GN2036 and *L. plantarum* GN104 were used to treat with the aquaculture water and aquaculture tail water. Ammonia nitrogen concentration of aquaculture water (a), nitrate concentration of aquaculture water (b), phosphate concentration of aquaculture tail water (c) and COD (d) indexes of aquaculture tail water were selected to evaluate the performance of each strain. DPI, days post-inoculation; COD, chemical oxygen demand.

## Conclusions and Discussion

This study introduced the innovative concept of DME and developed the DME application system, which was composed of a core hardware named DME intelligent fermentor (Fig. 1). We conducted fermentation tests on nearly 40 commonly used microbial agents, reaching a fermentation yield of 1∼10×10^8^ CFU/mL within 16∼48 hours, in the fields of planting, aquaculture and Silage (Table 2). Selected strains were then applied in fermentation trials, demonstrating excellent results in disease prevention and growth promotion for tomatoes, the improvement of fermented silage quality, and the improvement of water quality in aquaculture and aquaculture tail water (Fig. 2, 3; Table 3, 4). The “on-site” + “one-step” fermentation of the DME intelligent fermentor ingeniously avoided the issues of short shelf-life and unstable application effects of non-spore-forming functional microorganisms, while also bypassing the complicated sterile operation processes required in traditional large-scale or small-scale factory fermentation production. This model not only enhances production efficiency but also ensures the activity and functionality of the microorganisms, providing reliable support for practical applications.

Previously, the american company Pivot Bio developed a portable fermentation system for facultative anaerobic nitrogen-fixing bacteria, capable of achieving an expansion of 200 million bacteria per milliliter per day at room temperature (https://www.pivotbio.com/product-proven40-corn). However, this system was limited to anaerobic or facultative anaerobic bacteria, resulting in relatively limited application coverage. What’s more, the medium sterilization and aseptic filling were indispensable. Therefore, they through on-site fermentation solved the effectiveness and activity of functional strains while the complex aseptic filling progress and high transport cost still remains. In contrast, NBio accomplished the high pure fermentation for functional microorganisms bypassing the complex sterilization process by adopting directional selection pressure under a simple pasteurization process (85∼90°C, 15 min). In addition, the DME application system fully considered the fermentation needs of different types of strains by setting different aeration conditions to meet diverse fermentation requirements. This design makes the DME application system more flexible and efficient in handling various types of microorganisms, expanding its application scope and potential market.

Indigenous microorganisms, which form a natural group of microbial communities, existed in the soil and on the surfaces of all living organisms and play a role in biodegradation, nitrogen fixation, improving soil fertility, and plant growth promotion in agriculture (Firincă and Zamfir, 2023; Kumar and Gopal, 2015). The indigenous microorganisms can be dramatically different even the process of collection and isolation methods were similar and the application of indigenous microorganisms follows the rule below: where do they come from and where do they go to. It’s not realistic to establish standardized fermentation factories in each practical application scenario to achieve on-site production and application of indigenous microorganisms. Fortunately, the DME intelligent fermentor will bridge the gap between the isolation of functional indigenous microorganisms and their industrial application, facilitating the industrialization of these indigenous microorganisms with low cost and high efficiency. Additionally, its demonstrated the function of the functional microbial was carried out by its metabolites, including surfactant, surface polysaccharides, IAA, amino acid (Buescher and Margaritis, 2007; Femina Carolin et al., 2021; Jiang et al., 2020; Laird et al., 2020; Pérez-Burgos and Søgaard-Andersen, 2020), which had varying degree of lost during the formulation process or logistics transportation. Thus, the reduced content of bioactive of the commercial microbial agents was another main reason behind its poor application performance. Therefore, the application of the fermented microbial agents, with rich active metabolites, promising a desirable application performance by the DME intelligent fermentor and achieving the co-application of microbial agent and its metabolites.

In total, the DME intelligent fermentor not only exhibits superior application performance across multiple fields but also overcomes numerous challenges inherent in traditional fermentation methods through its innovative design and operational models. This positions the DME intelligent fermentor as a promising tool with substantial application prospects and market potential. The DME system was crucial for bridging the gap between basic research and industrial development of agricultural microorganisms, lowering the industrialization threshold for functional microorganisms, accelerating the application of functional strains, and promoting the healthy development of the agricultural microbial industry.

---

**Table S1.** Strains used in this study.

| Strains ID | Function | Species | Sources |
| --- | --- | --- | --- |
| GN125 | Plant growth, biocontrol | *Bacillus velezensis* | Green Nitrogen biotech Co., Ltd. |
| GN2036 | Water treatment | *Bacillus subtilis* | Green Nitrogen biotech Co., Ltd. |
| GN104 | Water treatment | *Lactobacillus plantarum* | Green Nitrogen biotech Co., Ltd. |
| GN251 | Feed fermentation | *Lactobacillus plantarum* | Green Nitrogen biotech Co., Ltd. |
| GN1367 | Feed fermentation | *Lactobacillus plantarum* | Green Nitrogen biotech Co., Ltd. |
| GN232 | Feed fermentation | *Lactobacillus plantarum* | Green Nitrogen biotech Co., Ltd. |
| GN1125 | Feed fermentation | *Lactobacillus plantarum* | Green Nitrogen biotech Co., Ltd. |
| GN1061 | Feed fermentation | *Lactobacillus buchneri* | Green Nitrogen biotech Co., Ltd. |
| GN1710 | Feed fermentation | *Lactobacillus casei* | Green Nitrogen biotech Co., Ltd. |
| GN456 | Feed fermentation | *Lactobacillus Faecalis* | Green Nitrogen biotech Co., Ltd. |
| RS | Pathogenesis | *Ralstonia solanacearum* | Green Nitrogen biotech Co., Ltd. |

**Table S2.**
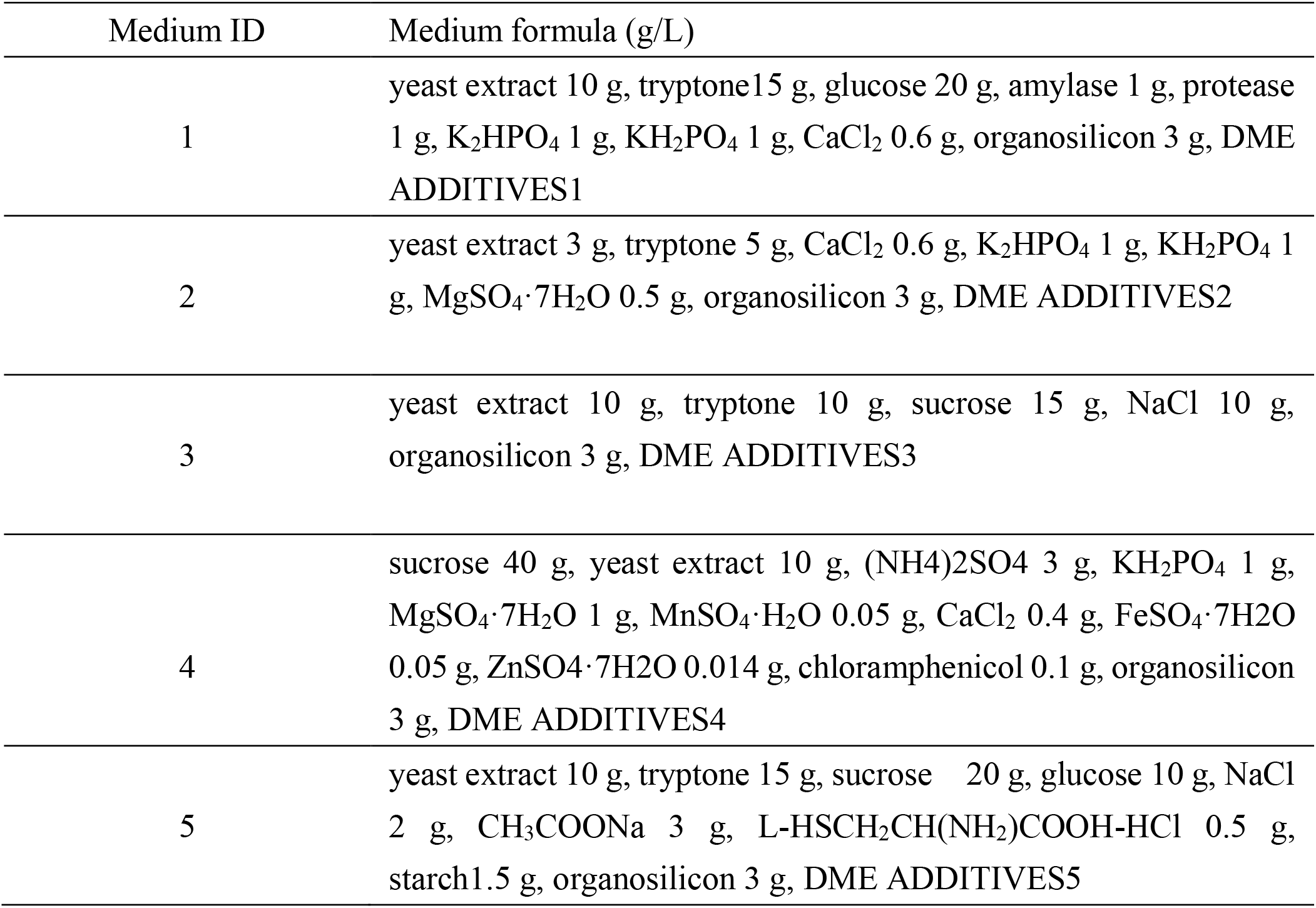
DME medium and its formula.

| Medium ID | Medium formula (g/L) |
| --- | --- |
| 1 | yeast extract 10 g, tryptone 15 g, glucose 20 g, amylase 1 g, protease 1 g, K<sub>2</sub>HPO<sub>4</sub> 1 g, KH<sub>2</sub>PO<sub>4</sub> 1 g, CaCl<sub>2</sub> 0.6 g, organosilicon 3 g, DME ADDITIVES1 |
| 2 | yeast extract 3 g, tryptone 5 g, CaCl<sub>2</sub> 0.6 g, K<sub>2</sub>HPO<sub>4</sub> 1 g, KH<sub>2</sub>PO<sub>4</sub> 1 g, MgSO<sub>4</sub> ·7H<sub>2</sub>O 0.5 g, organosilicon 3 g, DME ADDITIVES2 |
| 3 | yeast extract 10 g, tryptone 10 g, sucrose 15 g, NaCl 10 g, organosilicon 3 g, DME ADDITIVES3 |
| 4 | sucrose 40 g, yeast extract 10 g, (NH<sub>4</sub>)<sub>2</sub>SO<sub>4</sub> 3 g, KH<sub>2</sub>PO<sub>4</sub> 1 g, MgSO<sub>4</sub> ·7H<sub>2</sub>O 1 g, MnSO<sub>4</sub> ·H<sub>2</sub>O 0.05 g, CaCl<sub>2</sub> 0.4 g, FeSO<sub>4</sub> ·7H<sub>2</sub>O 0.05 g, ZnSO<sub>4</sub> ·7H<sub>2</sub>O 0.014 g, chloramphenicol 0.1 g, organosilicon 3 g, DME ADDITIVES4 |
| 5 | yeast extract 10 g, tryptone 15 g, sucrose 20 g, glucose 10 g, NaCl 2 g, CH<sub>3</sub>COONa 3 g, L-HSCH<sub>2</sub>CH(NH<sub>2</sub>)COOH-HCl 0.5 g, starch 1.5 g, organosilicon 3 g, DME ADDITIVES5 |

**Fig. S1.**
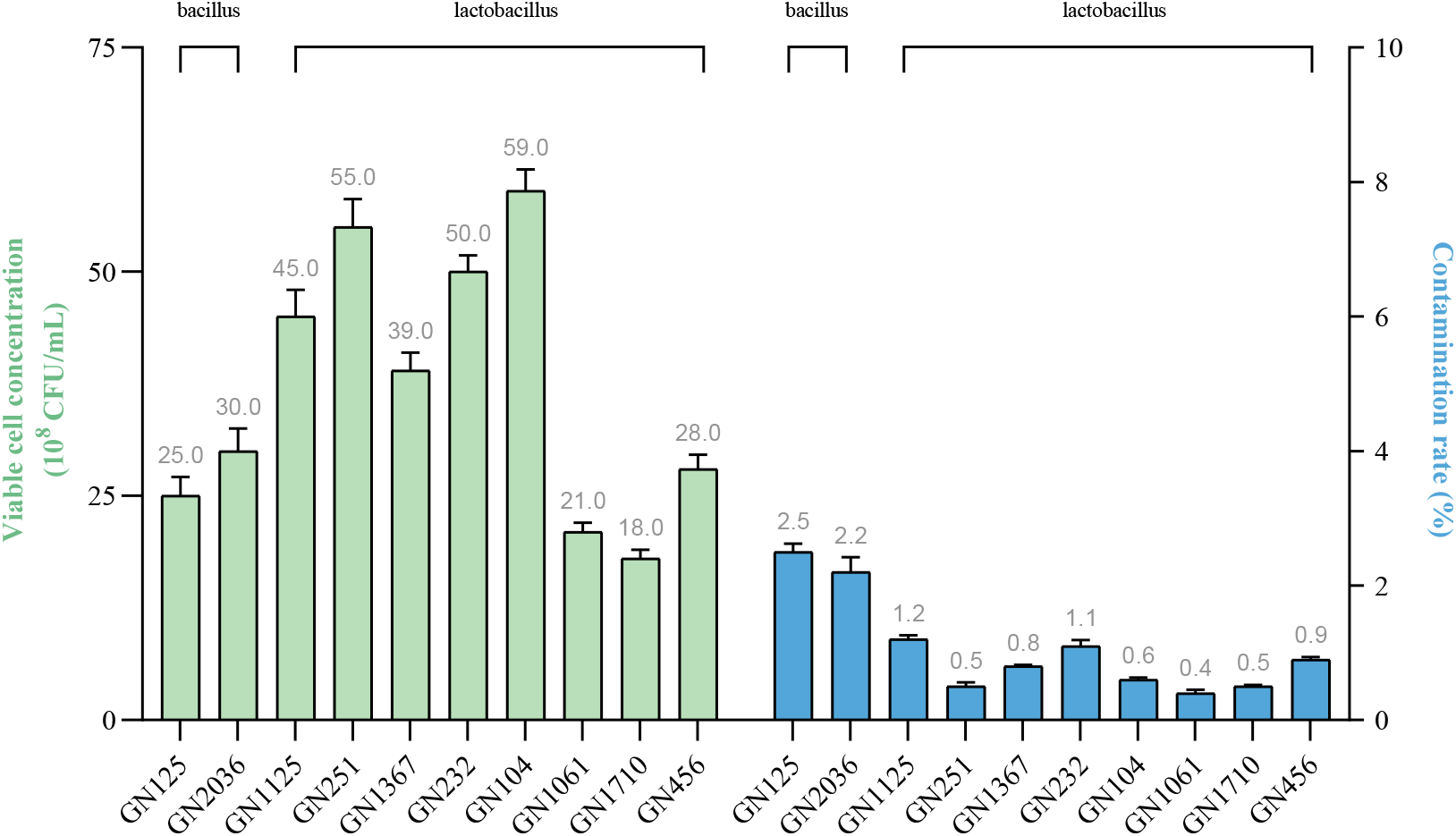
The used strains cultured by DME intelligent fermentor. The final cell concentration and contamination rate of used strains, including 2 bacillus strains and 8 lactobacillus strains. Error bars represents standard deviation (SD).

